# Standardizing a Protocol for Streamlined Synthesis and Characterization of Lipid Nanoparticles to Enable Preclinical Research and Education

**DOI:** 10.1101/2025.07.31.667476

**Authors:** Shilpi Agrawal, Abbey L. Stokes, Christopher E. Nelson

**Author notes:** Equal work contribution (Shilpi Agrawal, Abbey L. Stokes). **Address for correspondence** Christopher E. Nelson, Ph.D., Department of Biomedical Engineering, 120 John A. White Jr. Engineering Hall, University of Arkansas, Fayetteville, AR 72701.

## Abstract

Lipid nanoparticles (LNPs) have revolutionized nucleic acid delivery, enabled the first FDA-approved RNAi therapy (Onpattro), and accelerated the development of mRNA vaccines during the COVID-19 pandemic. The success of LNP-based vaccines demonstrated the potential of these nanoparticles for broader therapeutic applications. As interest in LNP-based therapies expands, there is an urgent need for low-cost, reproducible synthesis protocols that can be readily implemented across a wide range of research settings. Traditional methods for LNP synthesis, such as pipette and vortex mixing, can yield inconsistent results. In contrast, microfluidic mixing offers better control over LNP properties but requires expensive equipment. This financial barrier prevents many laboratories from accessing cutting-edge LNP technology, slowing the pace of preclinical research and limiting the exploration of its full therapeutic potential.

To address this, we developed a standardized protocol for microfluidic mixing using a syringe pump and a commercially available microfluidic chip. This approach offers a cost-effective and reproducible method for LNP synthesis. The protocol details the synthesis of LNPs, physical characterization via dynamic light scattering, encapsulation efficiency using the RiboGreen assay, and evaluation of *in vitro* transfection efficiency using the OneGlo assay, confocal microscopy, and a flow cytometer. We explored consistency through various experimental parameters, aiming to optimize the protocol and, importantly, tested user-to-user reproducibility with undergraduates without LNP experience with minimal supervision.

We found consistency among assembly conditions, including mRNA/lipid concentration, flow rate, dialysis time, ionizable lipid type, and chip reusability. We tested the broader applicability of the method with four ionizable lipids (DLin-MC3-DMA, LP01, C12-200, and SM-102). High encapsulation efficiency (96–100%) was maintained across tested concentrations, with slightly larger particle sizes at lower doses. Based on the flow cytometry analysis, a 7.5 µg dose was chosen for future use to conserve mRNA without sacrificing efficacy. Dialysis duration had minimal impact on encapsulation, but longer times increased particle size and reduced luminescence. The protocol consistently produced narrowly dispersed particles (PDI < 0.2), and equipment was reusable for up to six runs, potentially reducing the per-run cost. Even novice users achieved reproducible results, highlighting the protocol’s accessibility and potential to expand LNP research and applications.

## Introduction

Lipid nanoparticles (LNPs) are the most clinically advanced non-viral gene delivery vehicles, first approved by the FDA in 2018 to treat transthyretin-mediated amyloidosis by siRNA^1,2^. Moderna and Pfizer used LNPs to enable mRNA vaccines to address the COVID-19 pandemic^3,4^. The quick distribution to the public would not have been possible without prior clinical trials studying its safety *in vivo*. However, before LNPs entered clinical trials, they must undergo extensive combinatorial screening *in vitro* and *in vivo* to determine their mechanism of action and efficacy in drug delivery^5,6^. Decades of research have been spent optimizing the LNP components for various disease applications by targeting specific cell types and organs. LNPs are typically composed of 4 main components: an ionizable amino lipid, a helper phospholipid, cholesterol, and a PEGylated lipid^7^. Each of these components are responsible for characteristics of the LNP that contribute to overall functionality. The ionizable amino lipid is mainly responsible for the encapsulation of mRNA and endosomal escape^8,9^. The interaction between positively charged lipids at low pH during mixing and the negatively charged genetic material, alongside the hydrophobic effect, drives the lipids to enclose mRNA^7^. Helper lipids and cholesterol mostly contribute to the overall structure and stability of the LNPs^7^. The final component, PEG, improves the circulation time *in vivo* by reducing the rate of serum protein adsorption on the surface and targeting the LNP for destruction by phagocytes^10^. Biocompatibility, combined with the precise control of these physicochemical properties, has been crucial to the success of LNPs in human clinical trials, highlighting the importance of scalable and reproducible manufacturing approaches.

Current methods of manufacturing LNPs are distinct from traditional liposome formation by thin film hydration. These include 3 different formulation methods: pipette, vortex, and microfluidic mixing^11,12^. The limitations of pipette and vortex mixing include reproducibility due to increased human error, and the inability to scale up for large-scale production^13^. These methods are typically used in laboratory settings for small-scale production. Microfluidic mixing has shown promise for the large-scale production of nanoparticles for various applications, including preclinical and clinical studies^11,14^. Microfluidics mixing offers several advantages over traditional methods, such as batch synthesis, higher production rate, improved reproducibility, and greater control over particle size and distribution^15^. In the context of laboratory animal studies, the use of microfluidic mixing methods can improve control over size and encapsulation efficiency, which is essential for ensuring the safety and efficacy of experimental treatments^6,15,16^. In industry, large-scale mixing methods including T-junctions and confined impingement jet mixers are used to improve efficiency and reproducibility in the large-scale, GMP-compliant production of nanoparticle-based products.

To accelerate early-stage preclinical research and formulation screening of LNPs in diverse disease contexts, microfluidic mixing offers a reproducible and efficient method for LNP formulation. While large-scale production may employ T-junction or other scalable approaches, microfluidics remains a valuable tool for rapid testing and optimization in laboratory settings. However, the current limitations of this method include cost- and time-intensive nature of implementing this technique. For example, microfluidic instruments are expensive for early-stage research labs seeking preliminary data and grant funding. Microfluidic chips can be purchased off-the-shelf from various vendors, or they can be custom-made to meet specific research needs^16,17^. However, designing and fabricating custom microfluidic chips can be challenging and time-consuming, and may require specialized skills, and equipment. Purchasing off-the-shelf microfluidic chips may be a more convenient option for some researchers. Ultimately, the choice between buying off-the-shelf chips and making custom ones depends on the specific needs and resources of the researcher.

Here, we describe a relatively low-cost technique that incorporates microfluidic mixing using a syringe pump and microfluidic chip, with the system optimized to minimize dead volume and reduce material waste (**Fig. 1**). Rather than using the standard DLin-MC3-DMA^1,2,18^ formulation that was developed specifically for siRNA delivery by Onpattro, we used a 5-component lipid formulation. These 5-component lipid formulations containing 50% DOTAP were previously used *in vivo* to enhance lung targeting^11,19^. Therefore, we incorporated DOTAP^11,19^, a cationic lipid, to enhance encapsulation and transfection efficiency. This standardized protocol is relatively cost- and time-efficient and can be quickly integrated into most lab environments. To create the most efficient protocol, we investigated several factors involved in manufacturing and reproducibility. First, we evaluated LNP synthesis parameters to determine the most effective and cost-efficient conditions for our method. The parameters evaluated were the assembly concentration, total flow rate, dialysis time, and stability. We systematically varied the absolute concentrations of mRNA and lipids while keeping the total formulation volume constant, in order to identify the minimum input quantities required to achieve efficient encapsulation and *in vitro* transfection. Importantly, this experiment was not designed to evaluate dose-response effects or to optimize the N:P ratio, but rather to determine practical lower limits of reagents that still yield functional LNPs in a resource-efficient manner. This is critical for reducing reagent costs while maintaining transfection efficiency. Additionally, we stored these LNPs for 30 days at 4°C to determine their stability. Then, we optimized the total flow rate, testing 4, 8, and 12 mL/min, to evaluate its impact on LNP formation and uniformity. Regarding dialysis, some protocols recommend dialysis for 16 hours or overnight, while others reported successful results with as little as 1–2 hours^19–23^. We compared total dialysis times to assess whether shorter dialysis impacts LNP characteristics, aiming for time efficiency without compromising quality. The conditions found through these optimization steps were applied to 3 additional benchmark LNP formulations incorporating the 5-component lipid method^11,19^ alongside MC3:

**Figure 1.**
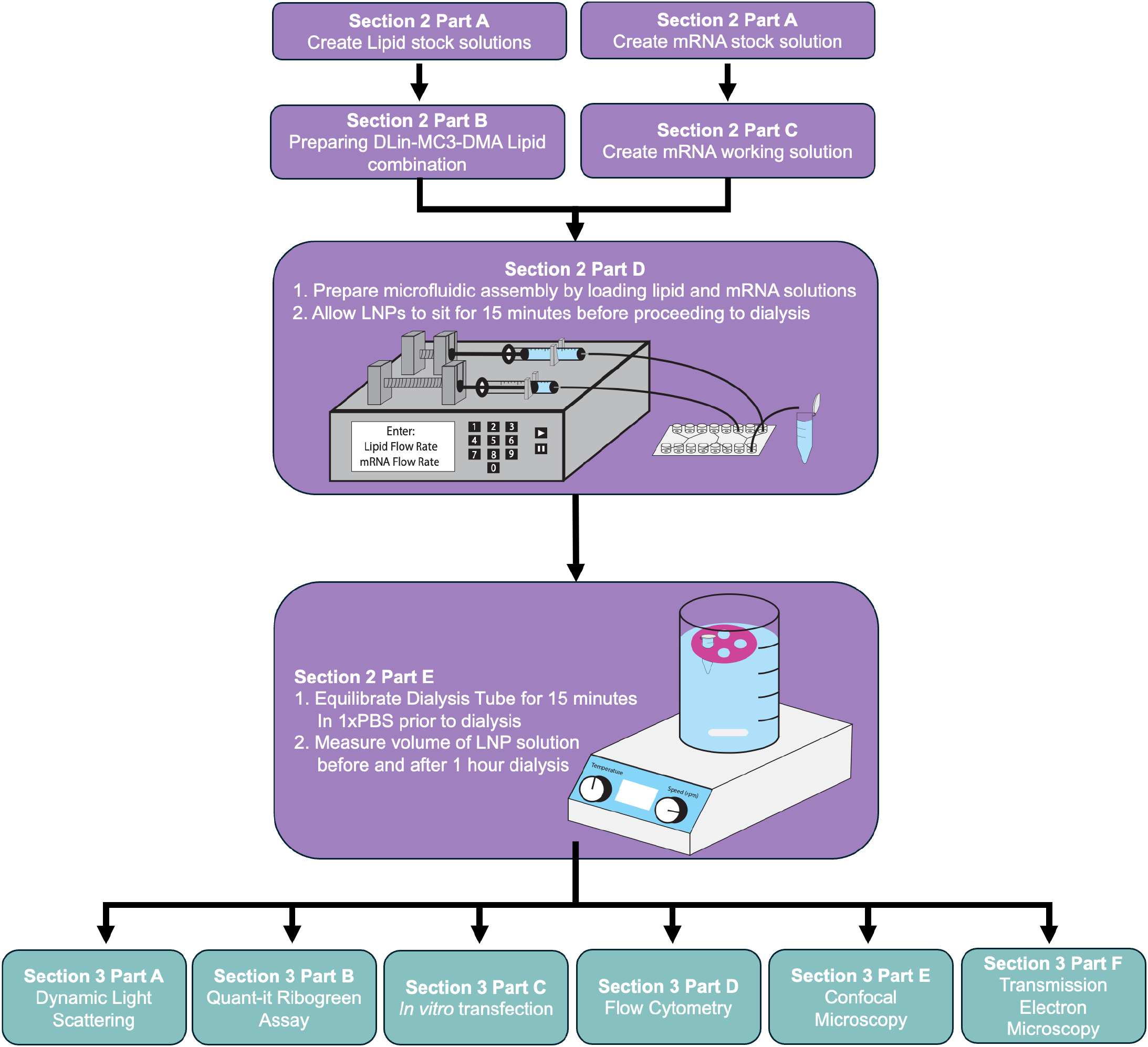
Overview of methods detailing the order of steps for LNP synthesis and characterization, along with the corresponding sections.

1. MC3:DSPC:Chol:DMG-PEG:DOTAP; 25:5:19.25:0.75:50^18^
2. LP01:DSPC:Chol:DMG-PEG:DOTAP; 22.5:4.5:22:1:50^24^
3. SM102:DSPC:Chol:DMG-PEG:DOTAP; 25:5:19.25:0.75:50^6^
4. C12-200: DOPE: Chol: DMG-PEG: DOTAP; 17.5:8:23.25:1.25:50^25^

Once we defined the ideal parameters to achieve successful LNP synthesis through our time and cost-efficient method, we evaluated microfluidic chip reusability. Currently, these channels are typically discarded after one run. However, from a cost and sustainability standpoint, it is important to determine whether repeated use impacts LNP quality-particularly in contexts such as teaching demonstrations or generating combinatorial LNP libraries where multiple runs are required. Therefore, we tested the effect of reusing the microfluidic chip and tubing on LNP self-assembly and particle uniformity. Through these iterative optimizations, we developed a standardized and reproducible protocol using a syringe pump setup that can be adapted for various lipid compositions. This cost-conscious approach enhances the feasibility of LNP-based research for a broader range of academic laboratories. To assess the accessibility and reproducibility of our protocol, we provided it to undergraduate students and evaluated whether they could successfully formulate LNPs with desirable physicochemical characteristics. This step helped us ensure that the method is not only robust and cost-effective but also simple enough to be implemented in educational and resource-limited lab settings.

## Materials and Methods

### Reagents

- LP01 (BP-Lipid 215, Broadpharm, cat. no. BP-26809)
- SM102 (Broadpharm, cat. no. BP-25499)
- C12-200 (Caymen Chemical, cat. no. 36699)
- DLin-MC3-DMA (MC3; MedKoo Biosciences, cat. no. 555308)
- Cholesterol, ≥99% (Avanti polar lipids, SKU no. 700100P)
- DMG-PEG (1,2-dimyristoyl-*rac*-glycero-3-methoxy(poly(ethylene glycol)) (Avanti Polar Lipids, SKU no. 880151P)
- DSPC (1,2-distearoyl-*sn*-glycero-3-phosphocholine) (Avanti Polar Lipids, SKU no. 850365P)
- DOPE (1,2-dioleoyl-sn-glycero-3-phosphoethanolamine) (Avanti Polar Lipids, SKU no. 850725P)
- DOTAP (1,2-dioleoyl-3trimethylammonium-propane) (Avanti Polar Lipids, SKU no. 890890P)
- NaCl (Sodium chloride) (Sigma Aldrich, cat. no. S9888)
- KCl (Potassium chloride) (Sigma Aldrich, cat. no. P3911)
- Na_2_HPO_4_ (Sodium phosphate dibasic) (Sigma Aldrich, cat. no. 567550)
- KH_2_PO_4_ (Potassium phosphate monobasic) (Sigma Aldrich, cat. no. P5655)
- Deionized water (facilitated by University of Arkansas)
- Absolute Ethanol (EtOH, 200 Proof; Fisher Scientific, cat. no. BP2818-500)
- DEPC treated water (EMD Millipore, cat. no. 693520)
- 10X PBS (Thermo Fisher Sci, cat. no. AM9625)
- Luciferase mRNA (GenScript, cat. no. SC2325)
- eGFP mRNA (GenScript, cat. no. SC2325)
- Sodium citrate dihydrate (Sigma Aldrich, cat. no. W302600)
- Citric acid monohydrate (Sigma Aldrich, cat. no. C1909
- PBS (1X), sterile filtered (Gennesse Scientific, cat. no. 25-508)
- Quant-it RiboGreen Assay Kit (Invitrogen, cat. no. R11490)
- Triton X-100 (Thermo Fisher Sci, cat. no. A16046.AE)
- Fetal Bovine Serum (FBS, Thermo Fisher, cat. no. A5670401)
- Dulbecco’s Modified Eagle’s Medium (DMEM, Genessee, cat. no. 25-500)
- Trypsin-EDTA (Fisher Scientific, cat. no. 25-200-056)
- Penicillin-Streptomycin (Gennesse Scientific, cat. no. 25-828)
- Lipofectamine 3000 (Thermo Fisher, cat. no. L3000001)
- HEK293T cells (ATCC, cat. no. CRL1573)
- One Glo Luciferase Assay system (Promega, cat. no. E6110)
- Uranyl acetate (Electron Microscopy Sciences, cat. no. 22400-4)

### Equipment

- pH meter (Ohaus, model no. ST3100)
- Analytical Balance (Gennesse Scientific, cat. no. 41-337)
- Syringe Pump (Chemyx, Model: Fusion 4000X)
- Microfluidic Chip (Lab Smith, SKU no. 10000076)
- Vortex-Genie 2 vortex mixer (Benchmark Sci, Model # WBB3113808)
- 1.7 mL microtubes (Gennesse Scientific, cat. no. 24-282)
- Glass vials – 5 mL, Amber (Uline, model no. S-25275A)
- 1 mL Luer Lock Syringe (VWR, cat. no. 76124-658)
- 3 mL Luer Lock Syringe (VWR, cat. no. 76124-660)
- Mini Luer to Luer adaptor (Lab smith, cat. no. 10000063)
- Connector tubing (Cole Parmer, cat. no. EW-50106-69)
- Long tubing and collection tubing (Lab Smith, cat. no. 10000033)
- Luer fluid connector (Lab Smith, cat. no. 10000080)
- Weighing Dishes (Gennesee Scientific, cat. no. 21-156)
- Parafilm (Gennesee Scientific, cat. no. 16-102)
- Pipettes and pipette stand (Eppendorf, cat. no. 2231001122)
- Pipette Tips (VWR, 1000 μL: 76175-386, 200 μL: 75841-126, 10 μL 75841-122)
- Weight to secure microfluidic chip
- Pur-a-lyzer Maxi Dialysis Kit (Sigma, SKU no. PURX35050)
- Floaters (Supplied via the Pur-a-lyzer Maxi Dialysis Kit)
- 1 or 2L beaker (VWR, cat. no. 76019-270)
- Magnetic stirrer (VWR, cat. no. 76447-052)
- Magnetic stirrer bar (Gennesse Scientific, cat. no. 27-530SL50)
- Aluminum Foil
- 37 °C incubator supplemented with 5% CO_2_ atmosphere (Eppendorf, cat. no. 671010015)
- Cell Sorter (Sony MA900)
- Biosafety cabinet level II (Labconco, cat. no. 76317-622)
- Transmission Electron Microscope (JEOL, cat. no. JEM-1011)
- Zetasizer Nano ZSP ZEN3600 (Malvern Instruments)
- Microplate reader (Biotek Synergy LX multimode plate reader)
- Vacuum pump (VWR, cat. no. 75870-734)
- Bead bath with temperature control (Lab Armor, cat. no. 10-876-005)
- 96-Well plate, black and flat bottom (Corning, cat. no. CLS3916)
- 96-Well Cell Culture Plates, Flat Bottom Wells, TC Treated (Gennesse Scientific, cat. no. 25-109)
- 35 mm glass bottom petri dish (MatTek Life Sciences, cat. no. P35G-1.5-10-C)
- Disposable Cuvettes (Amazon, cat. no. BrandTech 759071D)
- 40 µm EASYstrainer Cell Sieves (Greiner Bio-One, cat. no. 542140)
- Parafilm (Gennesee Scientific, cat. no. 16-102)
- Formvar/Carbon 300 Mesh, Cu (Electron Microscopy Sciences, cat. no. FCF300-Cu)
- Tweezers (Amazon, cat. no. 7-SA-SE)

### Procedure

#### Section 1: Buffers

1X PBS

- Dissolve 8g of NaCl, 0.2 g of KCl, 1.44 g of Na_2_HPO_4_, and 0.245 g of KH_2_PO_4_ in 800 mL of deionized water. Adjust the pH to 7.4 and set the final volume to 1000 mL. Alternatively, to make 1 L of 1X PBS from 10X PBS, add 100 mL of 10X PBS to 900 mL of deionized water.

Sodium citrate buffer (100 mM, pH 3)

- Dissolve 0.55 g of sodium citrate dihydrate and 3.80 g of citric acid monohydrate in 160 mL of deionized water. Adjust the pH to 3 and set the final volume to 200 mL.

#### Section 2: Synthesis of LNP

DLin-MC3-DMA Formulation Details (detailed calculations in **Supplementary File 1**)

- Molar Percentage Ratio - 50:10:38.5:1.5:100 (MC3:DSPC:Chol:DMG-PEG:DOTAP)
- Lipid: mRNA weight ratio - 40:1
- mRNA:Lipid volume ratio - 3:1

##### Section 2.A: Preparing lipid and mRNA stock solutions

Dissolve individual lipid components in 100% ethanol to prepare lipid stock solutions at the molar concentrations listed below. Each stock is made according to the calculated volumes based on the target molar concentrations for the formulation.

### Stock Concentrations

- DSPC (Molecular Weight: 790.19 g/mol) – 7.5 mM
- Cholesterol (Molecular Weight: 386.65 g/mol) – 7.5 mM
- DMG-PEG (Molecular Weight: 2509.20 g/mol) – 2.5 mM
- DOTAP (Molecular Weight: 698.54 g/mol) – 10 mM
- MC3 (Molecular Weight: 642.10 g/mol) – 15.5 mM

### Method

1. Put 5-20 mg of each lipid on a weighing boat, and calculate the volume of ethanol based on the required molar concentration (see above) in the stock solution in the formula given below:

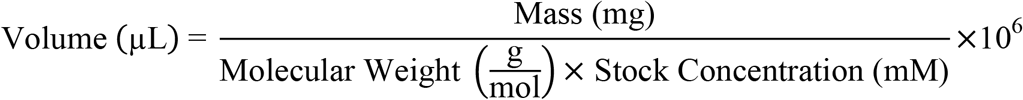

**Example:** 6.4 mg of DSPC is weighed. The required molar concentration needed is 7.5 mM. So, the volume of ethanol is calculated as follows:

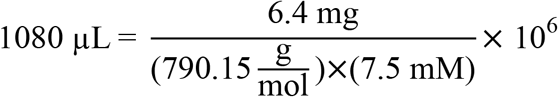
2. Vortex the mixture for 1–2 minutes until the lipid is completely dissolved. Aliquot the prepared lipid stock solutions into glass vials and store at −20°C until further use.
3. The mRNA stock solution is purchased at a concentration of 1 mg/mL. Upon receipt, it is immediately aliquoted into different 1.7 mL microtubes to avoid repeated freeze-thaw cycles.

#### Notes

- *Ensure all solutions are protected from light and moisture by storing in amber vials with tightly closed caps wrapped in parafilm*.
- *Use freshly prepared or appropriately stored stock solutions to avoid lipid degradation.*

##### Section 2.B: Preparing DLin-MC3-DMA Lipid combination

Combine:

1. 7.87 µL from 15.5 mM MC3 stock made in Part A
2. 3.27 µL from 7.5 mM DSPC stock made in Part A
3. 12.58 µL from 7.5 mM Cholesterol stock made in Part A
4. 1.47 µL from 2.5 mM DMG-PEG stock made in Part A
5. 24.51 µL from 10 mM DOTAP stock made in Part A
6. 265.30 µL of ethanol to a 1.7 mL microtube.

##### Section 2.C: Preparing mRNA working solutions for LNP formation

1. To prepare 7.5 µg of mRNA concentration in the LNP formulation, add 885.15 µL of DEPC-treated water to 99.23 µL of sodium citrate buffer and 7.88 µL of Luc/eGFP mRNA.

##### Section 2.D: LNP formation through Microfluidic Mixing

The microfluidic assembly should be prepared as shown in **Fig. 2A**. At this point, there are two 1.7 mL microtubes with: Luc mRNA (7.5 µg in 945 µL) and MC3 lipid combination (0.37 mM in 315 µL) as shown in **Fig. 2B**. The microfluid chip and connections are shown in **Fig. 2C**. For the following steps, the components used in the microfluidics assembly will be listed with their ID number **(Fig. 2D)**.

**Figure 2.**
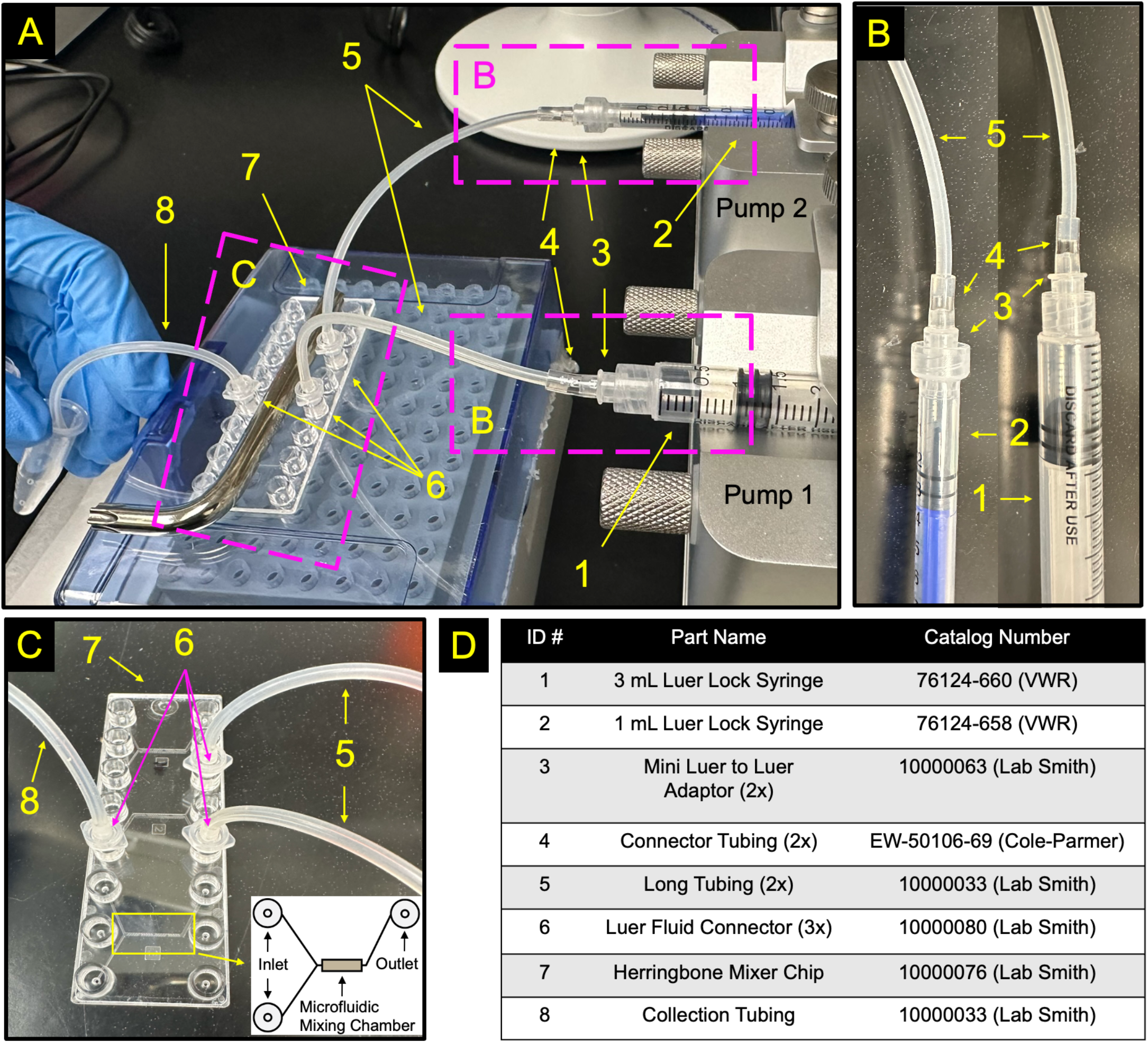
Schematic of microfluidic chip set-up. **A)** Picture of syringe pump with each part annotated in coordination with the table in the bottom right-hand corner. **B)** Close-up picture of syringe connections. Parts highlighted are outlined with purple boxes in A labeled with B. **C)** Close-up picture of the microfluidic chip connections separately. Parts highlighted are outlined with purple boxes in A labeled with C. A graphic provided on the lower right hand corner details one of three channel systems. **D)** Table coordinating the ID number with the part name used throughout the method section and the catalog number.

### Method

1. Cut two segments of approximately 1 cm of the connector tubing (#4).
2. Cut two segments of approximately 9-10 cm for long tubing (#5).
3. Connect the mini luer to luer adaptors (#3) to both luer lock syringes (#1 and #2) followed by the connector tubing (#4)
4. Put the long tubing (#5) inside the connector tubing (#4) (**Fig. 2B**).
5. Take up approximately 100 µL of any of the lipid stock in the 1 mL luer lock syringe and clear any air bubbles, if possible.
6. Take up approximately 300 µL of the mRNA in the 3 mL luer lock syringe and clear any air bubbles, if possible. *Note: No priming is necessary. However, for first-time users, running with deionized water first helps ensure proper assembly of the microfluidic chip, checks for tubing integrity and leaks, and verifies the syringe pump settings before proceeding with costly materials*.
7. Put 2 luer fluid connectors (#6) in the ports on the microfluidic chip (#7) for the first channel. This will act as an inlet for the lipid (ethanol) phase and the mRNA (aqueous) phase (**Fig. 2C**).
8. Put the two luer lock syringes in the syringe pump. When entering values into the syringe pump, the brand and values chosen were Norm Ject 1 mL for the lipids and Norm Ject 2.5 mL for the mRNA. The numbers programmed for each solution in the syringe pump are as follows:
  - Lipid: 0.3 mL and 3 mL/min
  - mRNA: 0.9 mL and 9 mL/min *Note: For more information on how to operate the syringe pump see* ***Fig. 3*** *caption. If using a different syringe pump or syringe size/brand, adjust the parameters accordingly based on the inner diameter and allowable flow rate. Most syringe pumps require entering the correct syringe dimensions or selecting from a preset list to ensure accurate flow rate and volume delivery*.
9. Put 1 luer fluid connector (#6) in the opposite port on the microfluidic chip (#7) for the first channel. Cut approximately 7 cm for collection tubing (#8). Connect it to the fluid connector (#6). This will act as the outlet channel to collect the LNPs (**Fig. 2C**).
10. Start the syringe pump. As the solutions mix through the channels, *let the first drop go on a Kimwipe prior to collecting the remaining* LNPs from the outlet tubing into a microtube.
11. Once completed, approximately 1.2 mL is collected in the microtube. (Slightly less than 1.2 mL of LNPs may be collected owing to the leakage and the loss of solutions in the syringes and connectors (typically 0.9-1 mL of LNPs is collected).
12. Let the final LNP solution sit at room temperature for 15 minutes and then transfer it to a dialysis tube.

#### Section 2.E: Dialysis

1. Add a magnetic stir bar to a 1L beaker filled with 1X PBS. Rehydrate and equilibrate the dialysis chamber by placing it in the PBS beaker with the help of a floater for 15 minutes. Cover with aluminum foil.
2. Measure the volume of the LNPs and mark it down in the Excel sheet as “volume after syringe pump”. (**Supplementary File 1**)
3. Transfer the formulated LNP to the dialysis tube and put it in the floater.
4. Immerse the floater with the dialysis tube in the 1L beaker.
5. Based on the dialysis time, put the magnetic stirrer at 4 °C or at room temperature. For overnight dialysis, put the magnetic stirrer at 4 °C, whereas for 1 or 2 hr dialysis, leave the magnetic stirrer at room temperature.
6. After the dialysis, transfer the solution into a microtube and record the “final volume”.
7. Use this LNP solution for further characterization.

**Figure 3.**
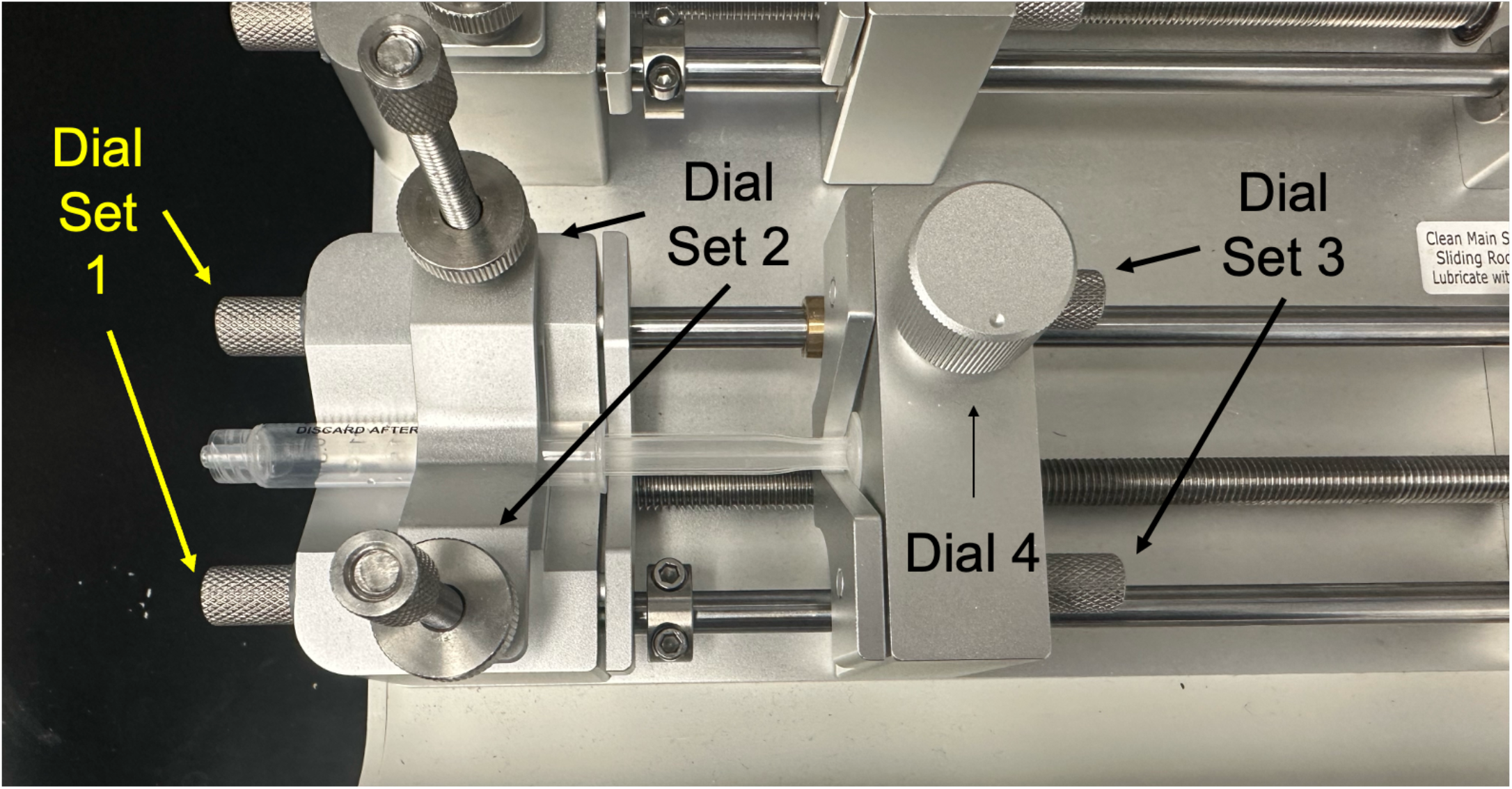
Close-up of a syringe loaded in pump 1 of syringe pump along with the required dials needed to secure syringe. Pump 2 dials are identical. Dial Set 1: Controls the opening where the flange of the syringe rests to restrict horizontal motion. Dial Set 2: Restricts syringe’s vertical motion Dial Set 3: Controls the opening where the end of the syringe’s plunger rests to ensure syringe pump is in full control of syringe’s motion. Dial 4: Turn counterclockwise (position depicted above) to adjust platform for syringe size. Turn clockwise (where dot is at the top) to run syringe pump.

##### Section 3: LNP Characterization

###### Section 3.A: Dynamic Light Scattering

1. Add 2900 µL of 1X PBS to the disposable cuvette.
2. Pipette 100 µL of the LNP formulation from section 2.E into the disposable cuvette and pipette mix.
3. Insert the cuvette into a Zetasizer Nano ZSP machine. Measure the size of the nanoparticle by using the following parameters:
  - For LNP: absorbance - 0.01, refractive index - 1.49
  - For PBS: viscosity – 0.8880 cP, refractive index – 1.330, dielectric constant – 79

###### Section 3.B: Quant-it Ribogreen RNA Assay

The ThermoFisher Scientific Quant-it RiboGreen RNA Assay Kit can be used with minor modifications to quantify both total and free mRNA in LNP formulations. The procedure consisted of two major parts: standard curve preparation and sample analysis.

### Method

Standard Curve Preparation

1. Prepare mRNA stock solution at 2 ng/µL using the same mRNA used in the LNP formulation.
2. Dilute the RiboGreen reagent 1:1000 with 1X PBS.
3. In a black 96-well plate:
  - Serially dilute the mRNA solution to generate a standard curve ranging from 2 ng/µL down to 0 ng/µL in 50 µL using PBS.
  - Add 50 µL of diluted RiboGreen reagent.
  - Cover the plate with foil and shake at 300 rpm for 5 minutes at room temperature.
4. Measure fluorescence at 485 nm excitation and 528 nm emission using a plate reader.
5. Add 50 µL of 0.5% Triton X-100 to each well, shake again at 300 rpm for 5 minutes, and re-measure the fluorescence.
6. Plot the standard curve and calculate the equation of the straight line (fluorescence vs. mRNA concentration):

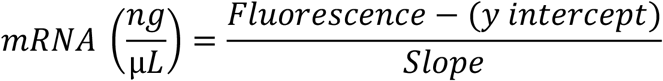

### Sample Analysis

1. For each sample:
  - Add 45 µL PBS, 5 µL of LNP sample, and 50 µL of diluted RiboGreen reagent to a well.
  - Cover and shake at 300 rpm for 5 minutes.
  - Measure initial fluorescence (to estimate free mRNA).
2. Add 50 µL of 0.5% Triton X-100 to each sample well (including standards), shake for 5 minutes, and re-measure the fluorescence (to estimate total mRNA after LNP disruption).

### Data Analysis

1. Total mRNA concentration (after Triton addition) is calculated using:

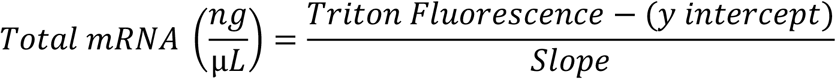
2. Free mRNA concentration (before Triton addition) is calculated similarly:

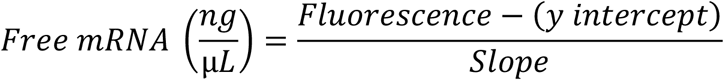
3. Encapsulation Efficiency (%) is determined by:

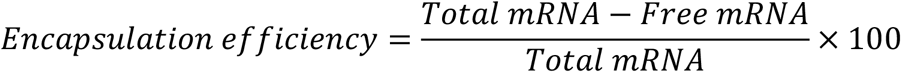

#### Section 3.C: In vitro transfection

##### Cell Dosing Calculations

Now calculate the volume of LNPs to be added to the cells based on the desired mass of mRNA. At this point, the following values should be known:

- Hypothetical mRNA (ng) – Section 2, Part C
- Hypothetical Volume (µL) – Section 2, Part D
- Volume After Syringe Pump (µL) – Section 2, Part E
- Final Volume (µL) – Section 2, Part E

##### Equations

1. Actual RNA (ng) - reflects the mRNA content proportionally recovered after syringe pump processing:

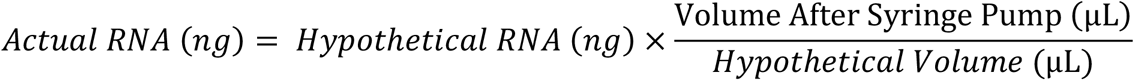
2. RNA encapsulated (ng) - Based on average encapsulation efficiency from Ribogreen assay in Section 3, Part C:

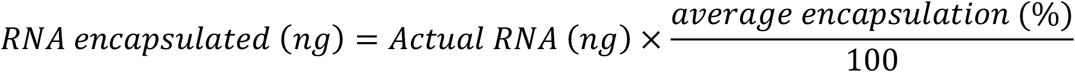
3. Encapsulated RNA concentration (ng/µL)

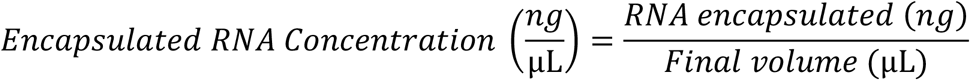
4. Volume to add to cells (µL) - Volume of LNP solution required to deliver a specific mRNA dose to cells:

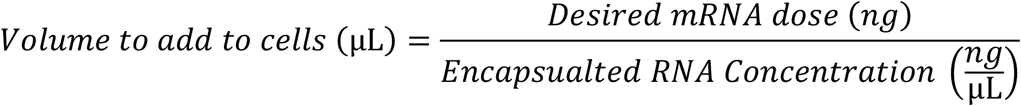

### Method

To test transfection efficiency, transfect HEK 293T cells, cultured in DMEM media supplemented 2with 10% FBS and 1% P/S. The positive control in our experiment is Lipofectamine 3000, and the untreated wells are considered as the negative control.

1. Seed 10,000 cells/well of a 96-well plate, and incubate the cells at 37°C in a humidified 5% CO^2^ incubator overnight
2. Calculate the volume of LNP to be added to each well based on the dosage (50 ng for the 96-well plates) using the equations above.
3. For Lipofectamine 3000 transfections, prepare mRNA-lipid complexes according to the manufacturer’s protocol.
4. For Luc mRNA, use the 96-well plates, and incubate the cells with LNP or Lipofectamine 3000 treatment for 16-18 hours. After 16-18 hours, measure luminescence with the One-Glo reagent following the manufacturer’s protocol. *Note: If GFP mRNA is used, seed 50,000 cells/well in a 24-well plate and transfect with 500 ng of LNPs. After 72 hrs, monitor GFP expression using flow cytometry*.

#### Section 3.D: Flow cytometry

Flow cytometry was performed using the Sony MA900 cell sorter (Sony Biotechnology). At approximately 72 hours post-treatment, detach all the cells by trypsinization and resuspend in PBS. Filter cell suspensions through 40 µm EASYstrainer cell sieves to remove aggregates. Each experiment includes three groups: blank/negative control (untreated cells), positive control (cells transfected using Lipofectamine 3000), and test samples (cells treated with LNPs). The gating strategy is depicted in **Supplementary Figure 1**. After data acquisition, discard all samples and analyze the data using FlowJo™ Software (BD Biosciences).

#### Section 3.E: Confocal microscopy

For confocal measurements, seed 300,000 HEK293T cells into a 35 mm glass bottom petri dish and transfect with 1000 ng of LNPs. At approximately 72 hours post-treatment, image the cells with an Olympus IX-83 inverted microscope using 20X magnification. A z-stack is taken and compiled using Fiji software.

#### Section 3.F: Transmission Electron Microscopy

Prepare LNP samples for TEM imaging with and without negative staining using carbon-coated copper grids.

1. Place a droplet (~5 µL) of the LNP sample on parafilm.
2. Place a TEM grid under the droplet (dull side up) and incubate for 2 minutes.
3. Then place the grid on a droplet of 4% uranyl acetate solution with the dull side facing down and incubate for 2 minutes.
4. Next, transfer the grid onto a droplet of ultrapure water (dull side down) and incubate for 2 minutes to remove excess stain.
5. Wick off excess liquid using a filter paper wedge placed at the edge of the grid.
6. Dry the grids overnight in labeled petri dishes and store in a desiccator before TEM imaging.

### Troubleshooting

When synthesizing LNPs, it is important to exercise caution when implementing certain strategies to address potential issues. Listed below are the strategies to address any potential problems that arise.

1. Initiating dialysis within 15 minutes of synthesis can help prevent LNP aggregation, but it is important to ensure that the buffer used for dialysis is appropriate for the specific lipid composition being used.
2. Thaw lipids at 45°C and vortex before adding them to the mixture. Ensure that mRNA and lipids are spun down separately at the end to ensure proper mixing. Avoid air bubbles in the syringe when loading lipid or mRNA. It is crucial to ensure that the tubing is flowing properly by testing it with water before use.

## Results

To streamline the development of LNP formulations, we optimized LNP synthesis parameters and cost-efficiency. The parameters focused on were the LNP assembly concentration, dialysis duration, total flow rate, and formulation stability over 30 days. Similar results were noted with three additional ionizable lipids and with GFP mRNA. We tested the reproducibility of the microfluidic system to lower the cost per run to increase the cost-efficiency of the LNP synthesis procedure. We also examined user-to-user variability with 7 independent users with limited LNP synthesis experience.

### Optimization of mRNA Concentration

LNPs were formulated using varying the concentration of the assembly by reducing the amount of mRNA (10 µg, 7.5 µg, 5 µg, and 2.5 µg), maintaining a constant lipid-to-mRNA weight ratio of 40:1. The mRNA volume of 900 µL and lipid volume of 300 µL remained unchanged. The ionizable lipid employed was MC3. Despite significant differences between a few groups in size and encapsulation efficiency, all LNPs showed favorable characteristics as found in literature^15,24,26^. Luciferase activity, assessed using the One-Glo assay, was highest in the 10 µg formulation, followed closely by the 7.5 µg sample (**Fig. 4A**). DLS measurements revealed well-controlled particle sizes of 82.3 nm (10 µg), 78.3 nm (7.5 µg), 82.3 nm (5 µg), and 101.02 nm (2.5 µg), with polydispersity index (PDI) values consistently below 0.2 across all samples (**Figs. 4B-C**). RiboGreen assay results indicated high encapsulation efficiencies (~98%) for all formulations (**Fig. 4D**). These results were verified with EGFP mRNA where 7.5 µg showed a comparable fluorescence intensity profile compared to 2.5 µg and 1 µg (**Supplementary Fig. 2**). Based on these results, 7.5 µg was selected as the optimal mRNA input for subsequent experiments, balancing performance and resource efficiency.

**Figure 4.**
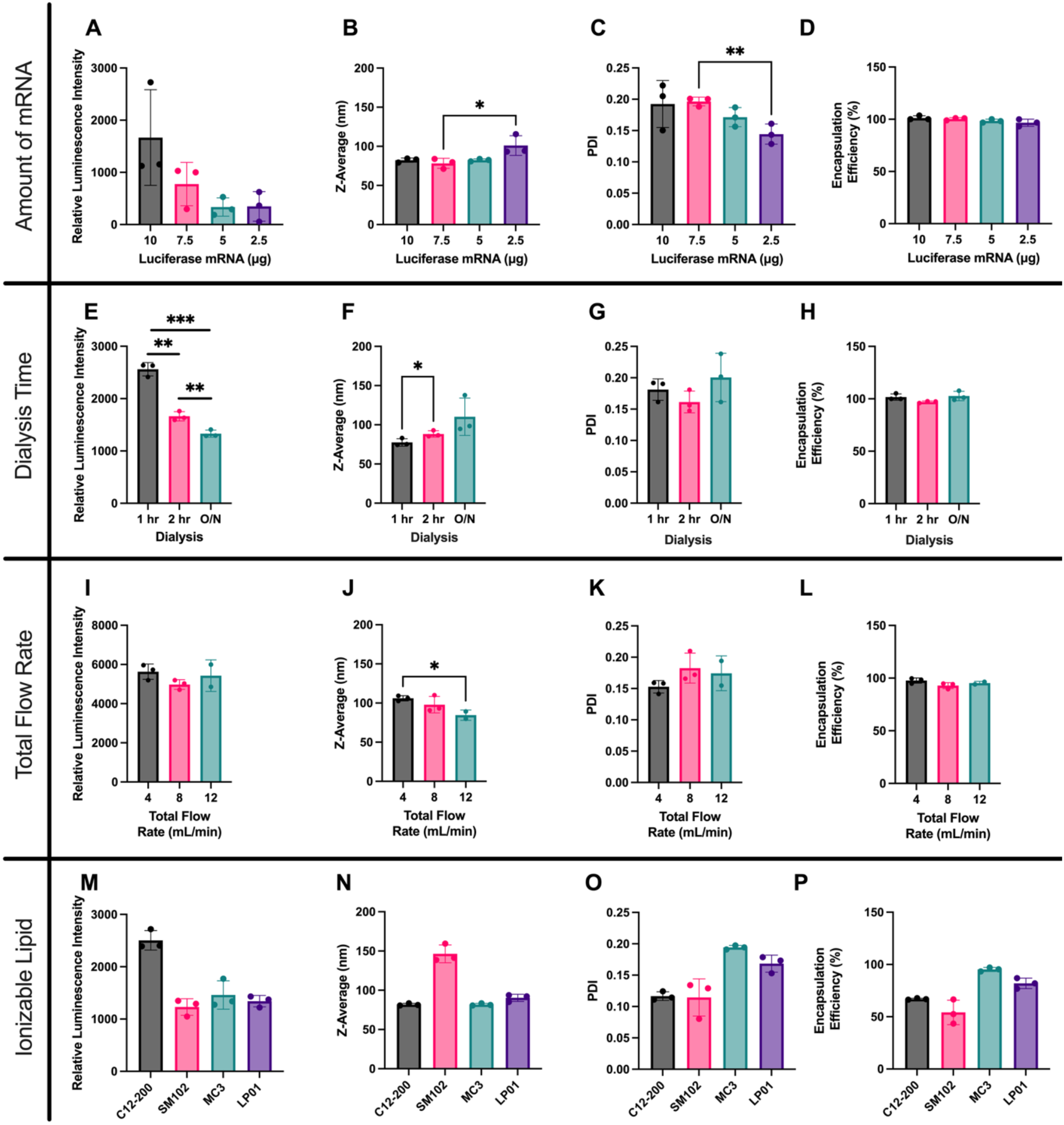
LNP characterization under varying experimental conditions. **(A–D)** LNPs synthesized with increasing amounts of luciferase mRNA in a fixed volume of 945 µL were characterized for (A) relative luminescence intensity, (B) particle size, (C) polydispersity index (PDI), and (D) encapsulation efficiency. **(E–H)** Effects of dialysis duration (1 hour, 2 hours, and overnight) on (E) luminescence, (F) size, (G) PDI, and (H) encapsulation. **(I–L)** LNPs produced at total flow rates of 4, 8, and 12 mL/min were assessed for (I) luminescence, (J) size, (K) PDI, and (L) encapsulation. **(M–P)** LNPs formulated with four lipid compositions—C12-200, SM-102, MC3, and LP01—were compared across (M) luminescence, (N) size, (O) PDI, and (P) encapsulation efficiency. Data are shown as mean ± SD. Statistical significance: p < 0.05 (**\***), p < 0.01 (**\*\***), p < 0.001 (**\*\*\***).

### Optimization of Dialysis Duration

Dialysis was conducted for 1 hour, 2 hours, and overnight (approximately 14-16 hours) to evaluate its effect on particle characteristics. Luminescence analysis revealed that the 1-hour dialysis condition produced the most robust transfection outcomes (**Fig. 4E**). The measured particle sizes were 77.5 nm (1 hr), 88.3 nm (2 hr), and 110.2 nm (overnight), with PDI values below 0.2 in all cases (**Figs. 4F-G**). RiboGreen assays confirmed encapsulation efficiencies exceeding 97% across all conditions (**Fig. 4H**). Therefore, a 1-hour dialysis time was selected for all subsequent formulations.

### Evaluation of Total Flow Rates

Three total flow rates—4 mL/min, 8 mL/min, and 12 mL/min—were assessed for their effect on the LNP properties. DLS results indicated comparable PDI values (<0.2) across all flow rates. Luciferase expression did not vary significantly among the conditions (**Fig. 4I**). Size measurements showed a significant decrease between 4 mL/min and 12 mL/min (**Figs. 4J-K**). This aligns with literature that has shown that a faster total flow rate results in a smaller LNP size^14,27^. However, encapsulation efficiency did not change with increasing flow rate: 97% (4 mL/min), 92% (8 mL/min), and 95% (12 mL/min) (**Fig. 4L**). Thus, 12 mL/min was selected for its operational efficiency in large-scale synthesis

### Stability Assessment Over 30 Days

Formulations prepared with varying mRNA amounts were characterized after 30 days of storage at 4°C. The 10, 7.5, 5, and 2.5 µg mRNA formulation all exhibited a large percent change in size. Then the 2.5 µg mRNA formulation exhibited the largest change in PDI, suggesting reduced colloidal stability (**Supplementary Fig. 3**). The instability was further demonstrated by large changes in encapsulation efficiency (Data not shown).

### Ionizable lipids

To evaluate the versatility of our LNP synthesis protocol, we tested four ionizable lipids: C12-200, SM102, MC3, and LP01^5,6,11,24,25^. SM102 and C12-200 produced larger nanoparticles (Z-average >130 nm), while MC3 and LP01 yielded smaller particles (~80–90 nm)(**Fig 4N)**. All formulations maintained low polydispersity indices (PDI < 0.2), indicating uniform particle populations (**Fig 4O**). Encapsulation efficiencies were highest for MC3 (~98%) and LP01, while SM102 and C12-200 showed lower efficiencies (~54% and ~66%, respectively) (**Fig. 4P**). Despite its lower encapsulation, C12-200 generated the highest luminescence, aligning with its established efficacy in in vitro applications (**Fig 4M**)^25^. These findings demonstrate that our protocol is adaptable across various ionizable lipids, including C12-200, SM102, MC3, and LP01, demonstrating consistent physicochemical properties and functional output., making it broadly applicable for educational and research settings.

### GFP mRNA Validation

We conducted a final validation experiment using 7.5 µg of GFP mRNA at a lipid:mRNA ratio of 40:1, with a total flow rate of 12 mL/min, 1-hour dialysis, and MC3 as the ionizable lipid. The resulting nanoparticles had an average hydrodynamic size of 80.84 nm and a PDI below 0.2 (**Supplementary Table 1**). Flow cytometry analysis demonstrated that 40.7% of the HEK293T cells were transfected with our MC3-LNP formulation (**Fig. 5A**). Additionally, LNP treated HEK293T cells were imaged using confocal microscopy (**Fig. 5B**). Transmission electron microscopy (TEM) confirmed uniform nanoparticle morphology with a size of ~50 nm (**Fig. 5C**). These results collectively confirm that our protocol yields efficient, reproducible, and cost-effective mRNA-loaded LNPs, suitable for both research and educational use.

**Figure 5.**
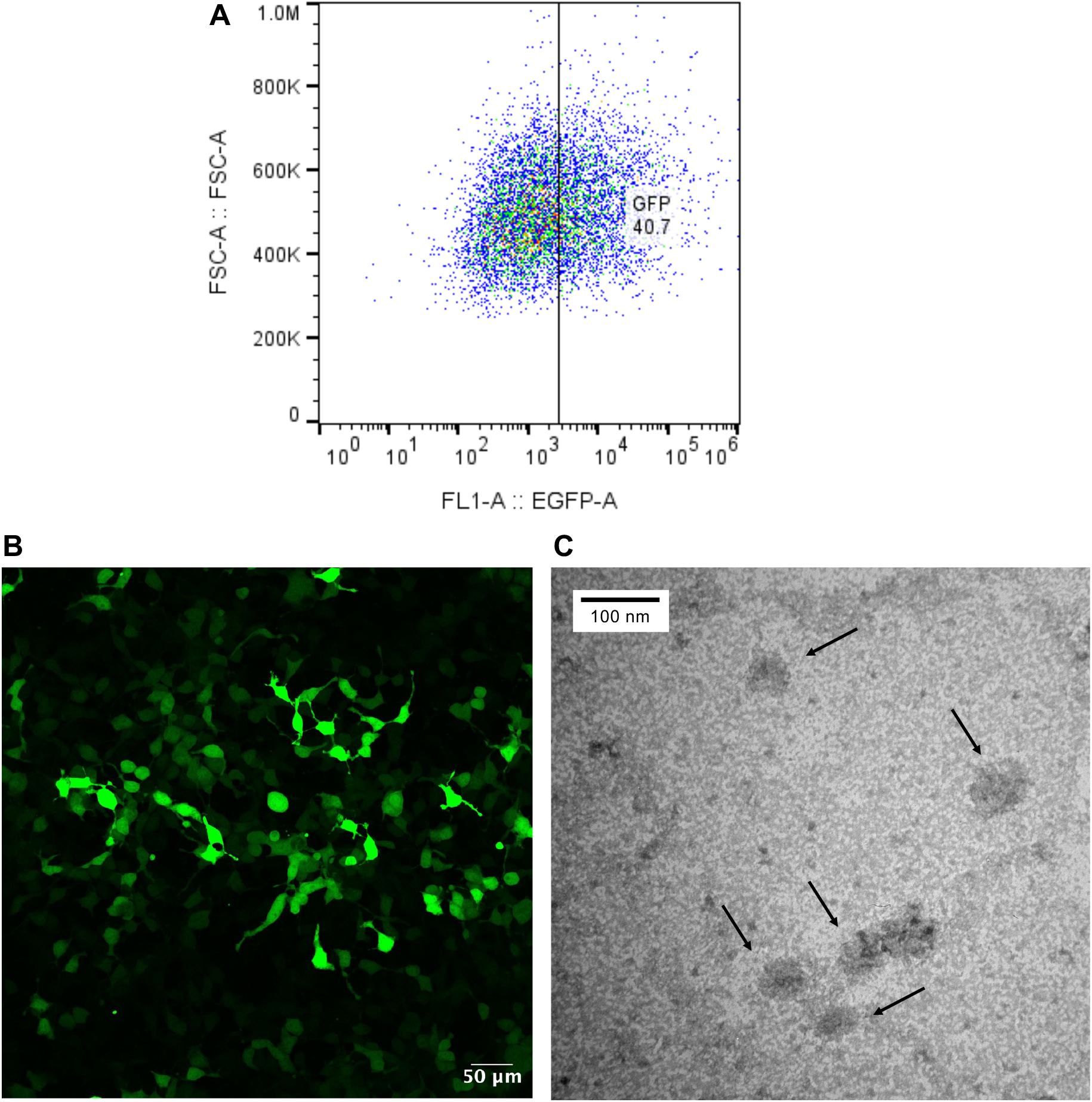
Results from final validation with GFP mRNA. **A)** EGFP fluorescence intensity following treatment with LNPs formulated using MC3, indicating successful mRNA delivery and expression. **B)** Image of HEK293T cells after LNP treatment using confocal microscopy. **C)** Image of LNP morphology using transmission electron microscopy.

### Chip Reusability

Since our aim is to enable undergraduate laboratories across the U.S. and globally to perform LNP synthesis with minimal resources, we tested whether the same microfluidic chip, along with its associated tubing and connectors, could be reused to reduce cost and material waste. To evaluate reusability, we conducted five consecutive LNP synthesis runs using the same herringbone mixer chip and monitored key formulation parameters, including particle size, PDI, encapsulation efficiency, and transfection efficiency. The percent change for each parameter was calculated relative to the first run (**Fig. 6A**). DLS measurements showed that particle size remained consistent across all runs, with less than 20% variation. Encapsulation efficiency remained stable, with changes within ±5%, and luminescence assays revealed negligible variability in transfection efficiency, suggesting preserved bioactivity of encapsulated mRNA. While the PDI showed a notable increase—over 50%—in the second run, this deviation may reflect experimental variability rather than a true decline in formulation quality. Subsequent runs showed moderate fluctuations, and all PDI values remained within the acceptable range (PDI < 0.2). These results demonstrate that the microfluidic chip and associated components can be reused for five additional runs without significantly compromising LNP quality, supporting a low-cost, accessible approach for teaching nanomedicine manufacturing. Additional runs could further decrease the cost per run but were not tested here.

**Figure 6.**
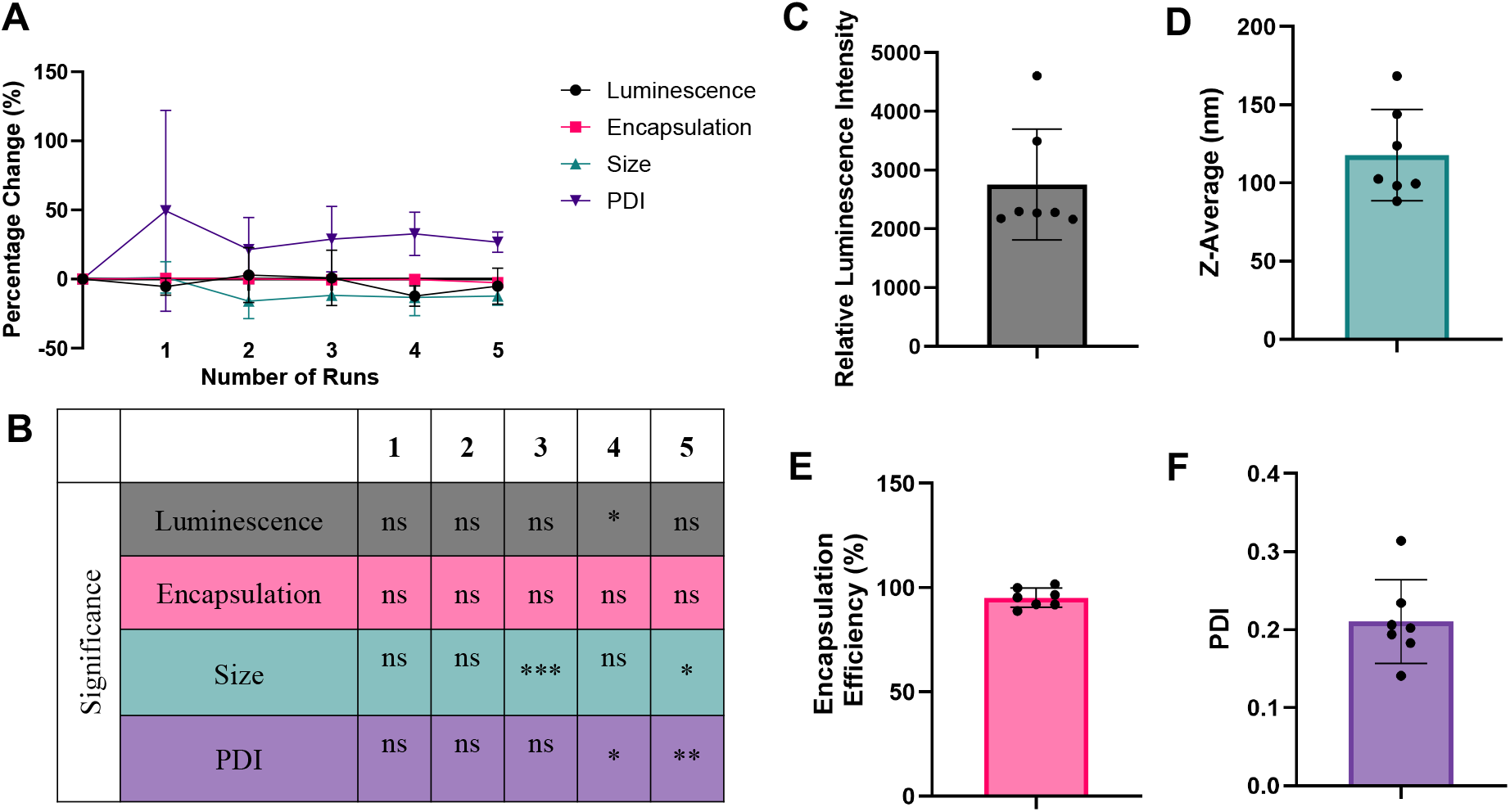
Results showcasing the application of our LNP synthesis protocol. **A)** Percent change of the relative luminescence intensity, encapsulation efficiency (%), size (nm), PDI after reuse of syringe pump tubing for an additional 5 runs. **B)** A table detailing any statistically different samples compared to the initial run (p<0.05 - *, p<0.01-**, p < 0.001 ***). **C-F)** Results from measuring relative luminescence intensity, encapsulation efficiency (%), size (nm), and PDI of 7 samples synthesized by undergraduates with minimal supervision.

### Reproducibility of LNP Synthesis by Undergraduate Students

To evaluate the reproducibility and robustness of our simplified LNP synthesis protocol, we provided a written version of the protocol—with no hands-on training or intervention—to seven undergraduate students, who independently synthesized MC3 LNPs. Despite their varied levels of laboratory experience, the students consistently produced LNPs with an average particle size that was 117.89 ± 27.12 nm, with a PDI of 0.21 ± 0.05, indicating nearly monodisperse populations. Encapsulation efficiencies were consistently high, averaging 95.46 ± 4.14%, and a relative luminescence intensity averaging 2755 ± 793, demonstrating functional delivery of mRNA (**Fig. 6C-F**). These findings confirm that our protocol is highly reproducible and can be implemented successfully by undergraduates, supporting its potential for broader adoption in teaching laboratories. Together, our results show that our protocol is effective, reproducible, and cost-efficient to allow its incorporation in academia and industry to allow expedited preclinical research of LNPs as a treatment method in various disease contexts.

### Timeline and cost analysis

LNP synthesis:
| **Sub process** | **Approximate time required** |
| --- | --- |
| Preparation of lipid stock solutions | 1-2 hours |
| Preparation of lipid and mRNA working solutions | 30 minutes to 1 hour |
| LNP synthesis | 30 minutes to 1 hour |
| LNP dialysis | 2-16 hours |

LNP characterization for each sample:
| **Sub process** | **Approximate time required** |
| --- | --- |
| Size and PDI measurement by DLS | 30 minutes to 1 hour |
| Encapsulation efficiency by RiboGreen assay | 20-30 minutes |
| In vitro delivery of the LNPs | 16-18 hours for luciferase activity and 48-72 hours for the GFP fluorescence |
| TEM measurement | 30 minutes to 1 hour (Prepare the negative stain sample 24 hours prior) |
| Confocal microscopy | 30 minutes to 1 hour (48-72 hours after in vitro transfection) |

Cost Analysis:
| **Sub process** | **Cost to Run 1 Sample** |
| --- | --- |
| LNP Synthesis | ~\$70 |
| LNP Synthesis (Reused 6 times) | ~\$25 |
| LNP Characterization | ~\$10 |
| **Initial Cost** |  |
| Syringe Pump | \$4399 |

Note: Amounts are approximations based on the method detailed in this protocol. Costs may vary depending on bulk ordering, characterization tests run, and more. These costs do not include the upfront cost of the syringe pump and other instrumentation needed throughout this protocol.

## Acknowledgements

We first would like to acknowledge the seven Nelson lab undergraduates who volunteered to make LNPs as part of this study. We would like to acknowledge the laboratory of Jin-Woo Kim for the use of the DLS instrument. We thank Erick Guererro and Professor Daniel Siegwart at the University of Texas Southwestern Medical Center for discussion and training on LNP preparation methods. All TEM imaging was performed after sample drying using standard operating procedures at the imaging core facility at the Institute for NanoScience and Engineering at the University of Arkansas. This work was supported by an NIH/NIBIB R00EB023979, NIH/NIGMS R35GM155433, NIH/NIGMS S10OD032338, ASGCT Career Development Award, University of Arkansas Chancellor’s Innovation Grant, and The Arkansas Bioscience Institute. CEN was supported by the 21st Century Chair in Biomedical Engineering. AS was supported by the Distinguished Doctoral Fellowship. SA was supported by a Women’s Giving Circle Grant.

## References

1. Coelho, T. et al. Safety and Efficacy of RNAi Therapy for Transthyretin Amyloidosis. N Engl J Med 369, 819–829 (2013).

2. Akinc, A. et al. The Onpattro story and the clinical translation of nanomedicines containing nucleic acid-based drugs. Nat. Nanotechnol. 14, 1084–1087 (2019).

3. Thomas, S. J. et al. Safety and Efficacy of the BNT162b2 mRNA Covid-19 Vaccine through 6 Months. N Engl J Med 385, 1761–1773 (2021).

4. Baden, L. R. et al. Efficacy and Safety of the mRNA-1273 SARS-CoV-2 Vaccine. N Engl J Med 384, 403–416 (2021).

5. Jayaraman, M. et al. Maximizing the Potency of siRNA Lipid Nanoparticles for Hepatic Gene Silencing In Vivo**. Angew. Chem. 124, 8657–8661 (2012).

6. Hassett, K. J. et al. Optimization of Lipid Nanoparticles for Intramuscular Administration of mRNA Vaccines. Molecular Therapy - Nucleic Acids 15, 1–11 (2019).

7. Eygeris, Y., Gupta, M., Kim, J. & Sahay, G. Chemistry of Lipid Nanoparticles for RNA Delivery. Acc. Chem. Res. 55, 2–12 (2022).

8. Semple, S. C. et al. Rational design of cationic lipids for siRNA delivery. Nat Biotechnol 28, 172–176 (2010).

9. Zelphati, O. & Szoka, F. C. Mechanism of oligonucleotide release from cationic liposomes. Proc. Natl. Acad. Sci. U.S.A. 93, 11493–11498 (1996).

10. Pozzi, D. et al. Effect of polyethyleneglycol (PEG) chain length on the bio–nano-interactions between PEGylated lipid nanoparticles and biological fluids: from nanostructure to uptake in cancer cells. Nanoscale 6, 2782 (2014).

11. Wang, X. et al. Preparation of selective organ-targeting (SORT) lipid nanoparticles (LNPs) using multiple technical methods for tissue-specific mRNA delivery. Nat Protoc 18, 265–291 (2023).

12. El-Mayta, R., Padilla, M. S., Billingsley, M. M., Han, X. & Mitchell, M. J. Testing the In Vitro and In Vivo Efficiency of mRNA-Lipid Nanoparticles Formulated by Microfluidic Mixing. JoVE 64810 (2023) doi:10.3791/64810.

13. Hirota, S., De Ilarduya, C. T., Barron, L. G. & Szoka, F. C. Simple Mixing Device to Reproducibly Prepare Cationic Lipid-DNA Complexes (Lipoplexes). BioTechniques 27, 286–290 (1999).

14. Zhigaltsev, I. V. et al. Bottom-Up Design and Synthesis of Limit Size Lipid Nanoparticle Systems with Aqueous and Triglyceride Cores Using Millisecond Microfluidic Mixing. Langmuir 28, 3633–3640 (2012).

15. Evers, M. J. W. et al. State-of-the-Art Design and Rapid-Mixing Production Techniques of Lipid Nanoparticles for Nucleic Acid Delivery. Small Methods 2, 1700375 (2018).

16. Belliveau, N. M. et al. Microfluidic Synthesis of Highly Potent Limit-size Lipid Nanoparticles for In Vivo Delivery of siRNA. Molecular Therapy - Nucleic Acids 1, e37 (2012).

17. Shepherd, S. J. et al. Scalable mRNA and siRNA Lipid Nanoparticle Production Using a Parallelized Microfluidic Device. Nano Lett. 21, 5671–5680 (2021).

18. Chander, N., Basha, G., Yan Cheng, M. H., Witzigmann, D. & Cullis, P. R. Lipid nanoparticle mRNA systems containing high levels of sphingomyelin engender higher protein expression in hepatic and extra-hepatic tissues. Molecular Therapy - Methods & Clinical Development 30, 235–245 (2023).

19. Cheng, Q. et al. Selective organ targeting (SORT) nanoparticles for tissue-specific mRNA delivery and CRISPR–Cas gene editing. Nat. Nanotechnol. 15, 313–320 (2020).

20. Whitehead, K. A. et al. Degradable lipid nanoparticles with predictable in vivo siRNA delivery activity. Nat Commun 5, 4277 (2014).

21. Thatte, A. S. et al. mRNA Lipid Nanoparticles for Ex Vivo Engineering of Immunosuppressive T Cells for Autoimmunity Therapies. Nano Lett. 23, 10179–10188 (2023).

22. Terada, T. et al. Characterization of Lipid Nanoparticles Containing Ionizable Cationic Lipids Using Design-of-Experiments Approach. Langmuir 37, 1120–1128 (2021).

23. Kimura, N. et al. Development of a Microfluidic-Based Post-Treatment Process for Size-Controlled Lipid Nanoparticles and Application to siRNA Delivery. ACS Appl. Mater. Interfaces 12, 34011–34020 (2020).

24. Finn, J. D. et al. A Single Administration of CRISPR/Cas9 Lipid Nanoparticles Achieves Robust and Persistent In Vivo Genome Editing. Cell Reports 22, 2227–2235 (2018).

25. Love, K. T. et al. Lipid-like materials for low-dose, in vivo gene silencing. Proc. Natl. Acad. Sci. U.S.A. 107, 1864–1869 (2010).

26. Chen, S. et al. Influence of particle size on the in vivo potency of lipid nanoparticle formulations of siRNA. Journal of Controlled Release 235, 236–244 (2016).

27. Kulkarni, J. A. et al. Rapid synthesis of lipid nanoparticles containing hydrophobic inorganic nanoparticles. Nanoscale 9, 13600–13609 (2017).

