## Supplemental Data for "Standardizing a Protocol for Streamlined Synthesis and Characterization of Lipid Nanoparticles to Enable Preclinical Research and Education"

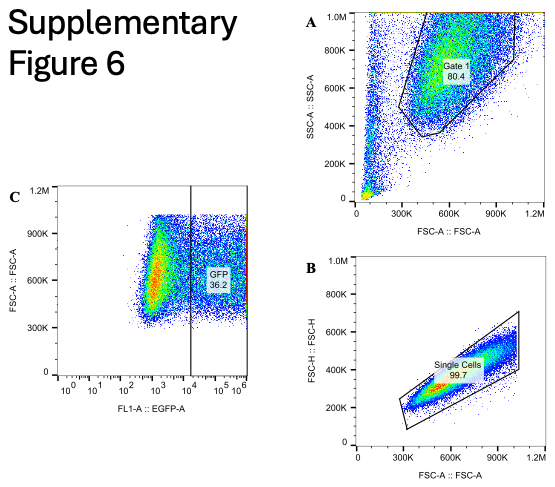

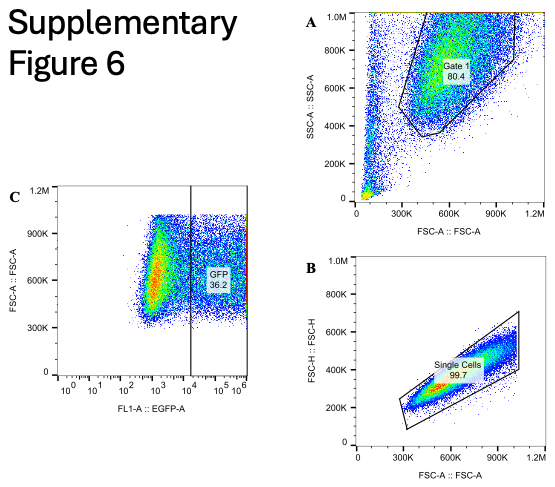

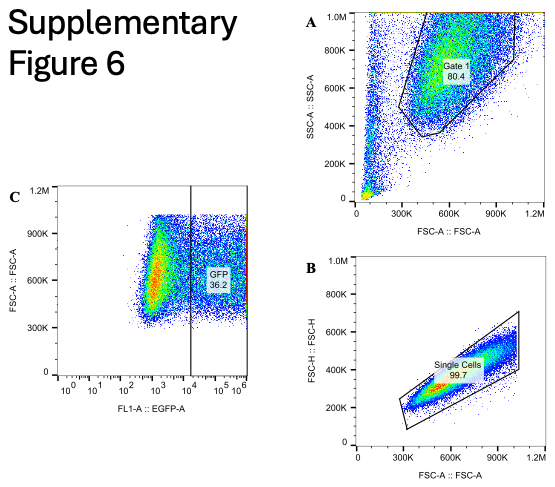

**Supplementary Figure 1**. Gating strategy used to analyze flow cytometry data of HEK293T cells after treatment by Lipofectamine, LNP, or untreated in Figure 6. A) FSC-A vs SSC-A isolates the cell population and eliminates cell debris. B ) FSC-A vs FSC-H isolates singlet cells from doublets. C) EGFP vs FSC-A was used to evaluate the percentage of EGFP% cells after treatment. Data was analyzed using the FlowJo^TM^ Software.

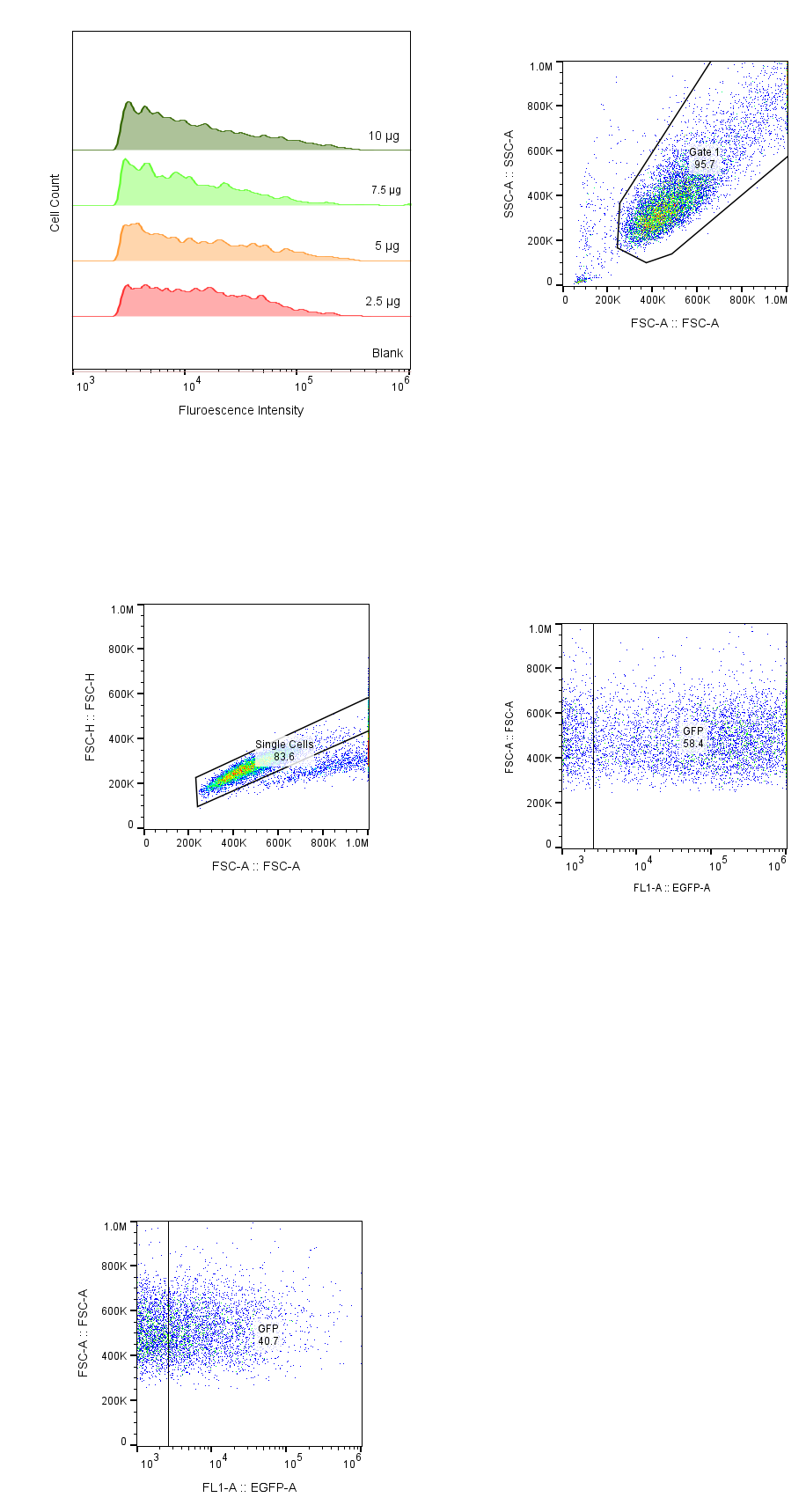
**Supplementary Figure 2**. Fluorescent intensity resulting from varying the amount of EGFP mRNA in the syringe pump system and transfecting HEK293T cells.

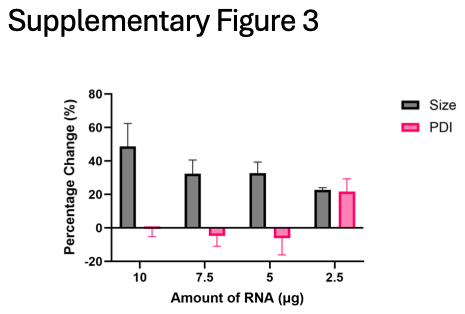
**Supplementary Figure 3**. Percent change in size or z-average (Black) and polydispersity index or PDI (Pink) after 4 weeks. Samples used were the amount of RNA samples (2.5 μg, 5 μg, 7.5 μg, 10 μg) depicted in Figure 1.

**Supplementary Table 1**. mRNA encapsulation efficiency, size or Z-average, and polydispersity index or PDI results when encapsulating GFP mRNA using the LNP parameters decided in the LNP optimization section.

| Size | PDI | Encapsulation |
| --- | --- | --- |
| 80.83 ± 3.25 nm | 0.198 ± 0.019 | 99.23 ± 2.66 % |

**Supplementary Table 2**. LP01, SM102, and C12-200 formulation parameters including: lipid composition, molar percentage ratio, lipid-mRNA weight ratio, and lipid-mRNA volume ratio. Concentrations of lipid stocks that are stored are: LP01 – 58.7 mM, SM102 – 140.8 mM, C12-200 – 88 mM.

| Size | PDI | Encapsulation |
| --- | --- | --- |
| 80.83 ± 3.25 nm | 0.198 ± 0.019 | 99.23 ± 2.66 % |

|  | LP01 | SM102 | C12-200 |
| --- | --- | --- | --- |
| Lipid Composition | LP01:DSPC:Chol:DMG-PEG:DOTAP | SM102:DSPC:Chol:DMG-PEG:DOTAP | C12-200: DOPE: Chol: DMG-PEG: DOTAP |
| Molar Percentage Ratio | 45: 9: 44: 2: 100 | 50:10:38.5:1.5:100 | 35:16:46.5:2.5:100 |
| Lipid:mRNA weight ratio | 10:1 | 10:1 | 10:1 |
| Lipid:mRNA volume ratio | 3:1 | 3:1 | 3:1 |
